# Development of a GFP Fluorescent Bacterial Biosensor for the Detection and Quantification of Silver and Copper Ions

**DOI:** 10.1101/296079

**Authors:** Adam R. Martinez, John R. Heil, Trevor C. Charles

## Abstract

Ionic silver is known to be an effective antimicrobial agent widely used in the cleaning and medical industries, however, there are several concerns regarding the release of silver pollutants into the environment. Presented here are two engineered bacterial biosensors for the detection and quantification of silver. The biosensors contain a silver resistance operon and a GFP gene that is strictly regulated through silver activated regulatory regions that control expression of the *sil* operons. The two biosensors are responsive to a wide range of silver ion concentrations, and a correlation between silver and GFP signal is seen at select concentration ranges. The biosensors were shown to detect silver ions released from silver nanoparticles, and have the potential to become a method for monitoring ion release rates of different nanoparticles. Interestingly, the close homology of the silver resistance and copper resistance genes allowed for the biosensor to also be responsive to copper ions, implying that copper ions activate silver resistance. Further development of this biosensor could lead to commercial applications for environmental monitoring.

**Importance:** Ionic silver is known to have many harmful environmental effects. Silver pollutants have been found in various environmental settings such as natural waterways and tailings from mining operations, raising concern. In addition, persistent exposure to silver in medical and environmental settings has led to the development of silver resistant bacteria, many of which are also resistant to a wide range of antibiotics. Some of these have the potential to develop into human pathogens. It then becomes important to have standardized methods for detecting and monitoring silver concentrations in various environments so that appropriate measures can be taken to prevent further silver ion release. This research shows that bacterial biosensors engineered to detect and quantify silver ions can be developed as effective alternatives to traditional analytical techniques. Further development of such biosensors could result in a commercial system for short and long term environmental monitoring, which is important as products containing silver and other heavy metals become increasingly popular.

## Introduction

Ionic silver is an effective antimicrobial agent widely used in the cleaning and medical industries (1, 2). However, this heavy metal has burdened the environment through its release in large quantities, and persistence as hazardous waste (3, 4). Silver ions and silver nanoparticles have repeatedly been reported in natural waterways and mining areas and are known to affect microbial communities, plants, and even animals (5).

There have been several efforts to develop methods for detecting and quantifying the extent of silver pollution within an environmental sample. Guo *et al.* developed a silver nanoparticle detection and quantification method by integrating a filtration technique into surface-enhanced Raman spectroscopy (SERS) (6). Zucker and Daniel developed a wide-field microscopy method for detecting silver and titanium nanoparticles (7). Stuart *et al.* demonstrated that particle-impact voltammetry (an electrochemical method) shows promise for detecting environmental nano-particle pollution (8). However, these methods require advanced spectroscopy methods, are expensive and not very portable or accessible when doing large scale environmental fieldwork. Thus, other methods for detecting and quantifying silver would be beneficial.

Several bacterial based biosensor systems have been engineered for the detection of various metal pollutants in environmental samples (9). Ravikumar *et al.* (10) developed a bacterial biosensor for zinc and copper with a detection limit of 16 μM and 26 μM respectively by utilizing CusRS and ZraRS two-component systems and a reporter gene, and have suggested a similar approach for detection of silver (11). Selifonova *et al.* (12) developed tree bioluminescent sensors for the detection of bioavailable Hg(II) in the environment with the highest sensitivity at 25 nM. Lastly, Liao *et al.* (13) developed and tested a GFP-based bacterial biosensor for the detection of bioavailable arsenic in contaminated groundwater samples. These sensors are becoming increasingly more applicable as alternatives to traditional analytical methods. An advantage of the use of biosensors is that contaminants are able to be quantified relative to the concentrations exposed to the biosensor, whereas traditional methods are relative to the extraction technique that is used for the analysis (14).

One approach to the development of whole-cell biosensors is to apply recombinant DNA technology to construct a plasmid vector where a regulated promoter drives expression of a reporter gene. The most effective promoters for environmental analysis are found in bacteria that are able to survive under conditions of extreme contamination by heavy metals (14). The ability to survive is sometimes mediated by a precisely regulated resistance system (1, 12). The design and construction of new biosensors could incorporate aspects of such systems.

The persistent exposure of silver in medical and environmental settings has led to the development of silver resistant bacteria (2). Most notable is a clinically isolated Ag(I)-resistant pathogenic *Salmonella*, from which the plasmid pMG101 was isolated in 1975 (15). The resistance mechanism (*sil* operon), has been characterized and is comprised of nine ORFs, two of which (*silR* and *silS*) encode a two-component silver sensor and regulator unit, and the other seven are structural genes (*silE*, *silC*, *silF*, *silB*, *silA*, *ORF105* and *silP*) that generally function by removing silver ions from the cytoplasm by membrane bound efflux pumps (15). This operon has been shown to convey resistance to 0.6 mM AgNO_3_, and is thereby a potentially effective mechanism for development of a biosensor to sense between 0 mM and 0.6 mM.

The two-component sensor and regulator unit *silR* and *silS* is able to detect available silver ions that are external to the bacterial cell (16, 17). External detection mechanisms, as opposed to internal detection mechanisms, are more accurate in quantifying heavy metal pollutants since factors like heavy metal uptake, efflux, and complexation do not need to be accounted for (16).

Lastly, for biosensors to be specific for the detection of a particular pollutant, the promoter which activates the reporter gene should ideally only become active in the presence of that specific pollutant. In the case of this mechanism, it has been demonstrated that copper ions activate the production of the resistance mechanism through *cusRS*, which is a copper-responsive, two-component sensor and regulator system. Since *cusRS* and *silRS* systems are assumed to function in similar manner, it was suggested that the silver ions could also activate the silver resistance mechanism through *silRS*.

One drawback of whole-cell biosensors is that they tend to only detect available ions in a media or environmental sample. If there happen to be various halides which can bind with the heavy metal in the media, the biosensor may not account for those ions. Though, for certain applications, only detection of available heavy metal ions is required (18).

Presented here are two GFP fluorescence based bacterial biosensors for silver. The two component silver sensor and regulator detection mechanism from the *sil* operon is coupled to green fluorescent protein expression. The sensor is the membrane bound protein (SilS) from *E. coli* J53(pMG101) which signals the activation of the resistance mechanism after it detects silver ions. This protein then activates the secondary messenger protein SilR by transphosphorylation, which then becomes a DNA binding activator (1). The promoter activated by phosphorylated SilR (of the *silE* and *silABC* genes) was cloned upstream of a promoterless *gfp* to create a transcriptional fusion, thus creating the plasmids pRADEK.1 and pRADEK.2, respectively. Both plasmids allowed for the detection of silver through GFP fluorescence. Such biosensors can also confirm and provide valuable insight into the effects of silver ions on transcription regulation of the *sil* operon. The function of *silR* and *silS* acting in a two component activation system is hypothesized due to these genes’ close homology to the copper resistance mechanism genes (19). This similarity raises the question of whether copper can activate silver resistance and *vice versa*. A correlation between GFP signal and silver ion concentration will not only provide a method for quantifying the ions, but perhaps further confirm suspected methods of *sil* operon regulation.

## Results

### Construction of two biosensor variants that are responsive to silver ions

Biosensing plasmids were constructed by fusing a mutGFP gene (see Materials and Methods) with silver regulated promoters on pMG101. Two biosensing plasmids were made, pRADEK.1 (containing *silE* promoter) and pRADEK.2 (containing *silABC* promoter). These constructs were then transformed into *E. coli* J53(pMG101) to create the bacterial biosensor strains. Both biosensor variants, RADEK.1 and RADEK.2, were able to produce a GFP signal in various silver concentrations that was statistically greater (p < 0.05) than signals produced by the biosensor with no silver present. Both RADEK.1 and RADEK.2 were able to detect concentrations of 0.04 mM to 5 mM in LB, and 0.04 mM to 0.31 mM in modified LB (LB with no sodium chloride), respectively (Figure 1). The different detection ranges observed between media (LB and modified LB) is expected since sodium chloride is known to form a AgCl precipitate which would decrease the availability of free silver ions. A lower amount of available silver ions in the media could assist the biosensor cells survival at higher absolute silver concentrations, resulting in a greater detection range, as was observed.

**Figure 1:**
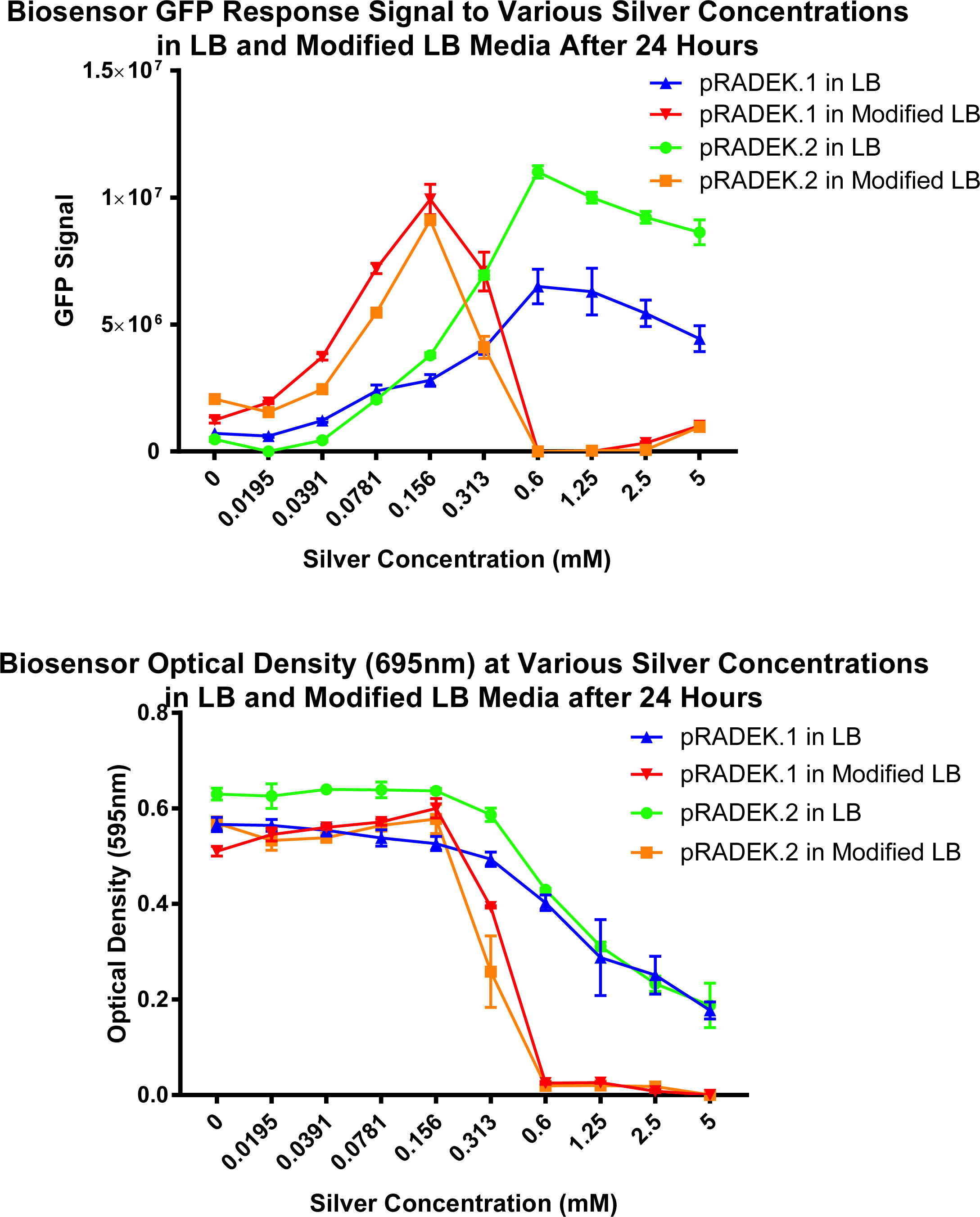
Line graph showing the optical density and GFP signal of the biosensors RADEK.1 and RADEK.2 in LB and modified LB at various silver nitrate concentrations.

### Correlation between GFP signal and silver ion concentration is detected with the biosensor variants

A correlation between GFP signal and silver ion concentration is detected with the biosensor variants. The correlation range varies depending on the media, and in specific segments of the detection range. Both RADEK.1 and RADEK.2 exhibited a strong correlation (R squared value above 0.96) in modified LB and LB (Figure 2). At concentrations above these ranges, the optical density of the biosensor decreased, resulting in a loss of correlation between silver and GFP. The different correlation ranges seen with different media (LB and modified LB) is expected, as AgCl precipitate influences the biosensor grow at higher silver concentrations.

**Figure 2:**
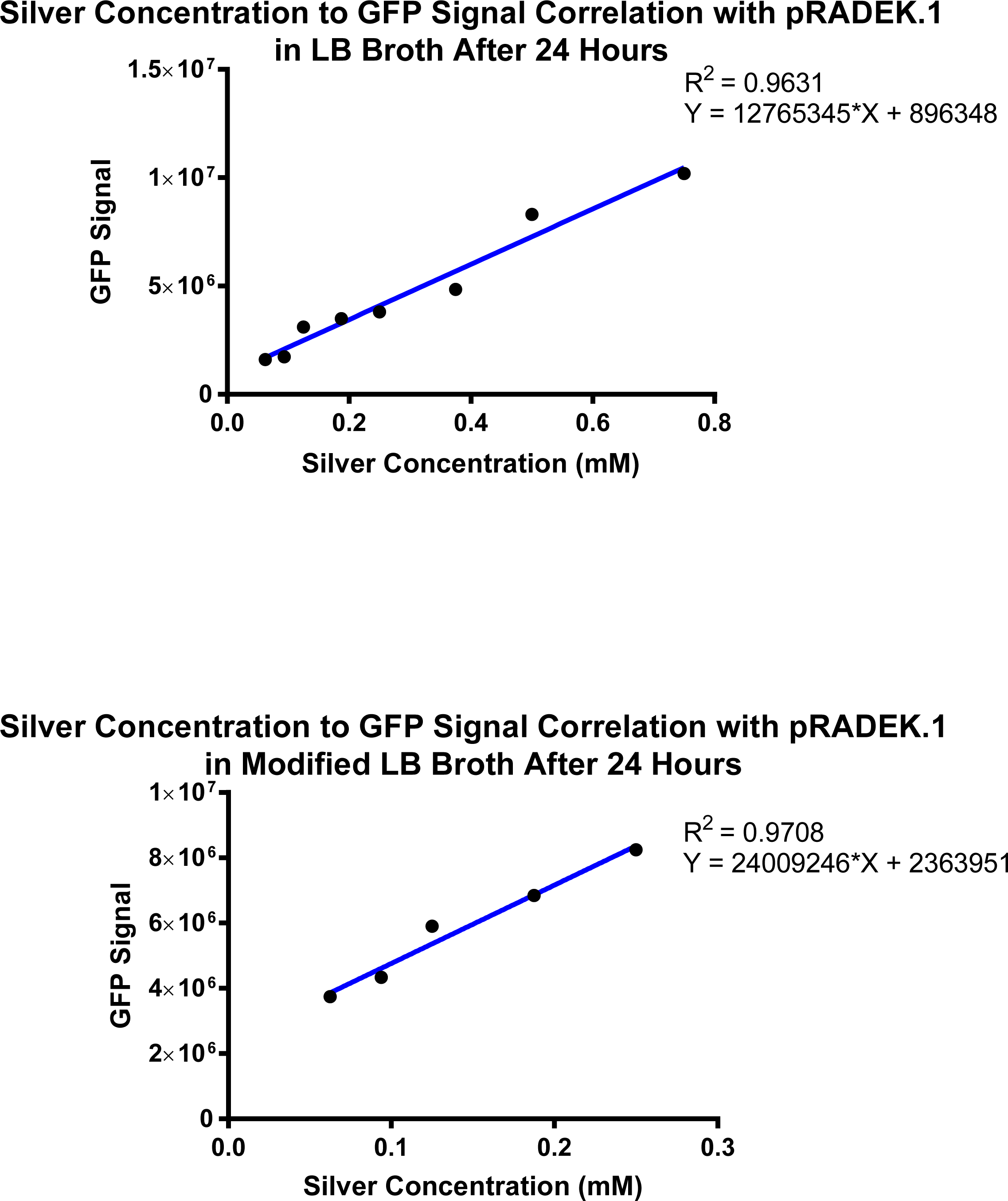
Correlation between silver concentration and GFP signal with biosensor RADEK.1 in LB and modified LB media. Linear equation and R squared value for each correlation are presented.

### Biosensor variants are able to detect silver nanoparticles

Two types of research grade nanoparticles (MR266 and PT163), were tested with both biosensors RADEK.1 and RADEK.2 starting at 0.1 mM and two-fold dilutions to 0.01 mM. These nanoparticles were described to be fairly stable by the manufacturers, meaning only 1% of silver ions would be in solution after synthesis, however, this can change depending on the oxidative environment. The GFP signal in both biosensor variants only increased by a significantly greater amount (p < 0.05) relative to samples containing no silver, at 0.1 mM concentration (data not presented). Similar to silver ions, at lower concentrations (between 0.05 mM and 0.01 mM) the signal decreased from samples containing no silver, again, likely due to noise within the sensing system. The optical density (595 nm) of the biosensor remained consistent throughout the different silver nanoparticle concentrations, thus, did not disrupt the accuracy of the GFP signal measurements.

### Biosensor variants are responsive to copper ions

The pMG101 plasmid is known to contain a copper resistance mechanism. Similar to silver ions, copper ions are able to be detected by the two biosensor variants in modified LB broth from concentrations 0.04 mM to 0.31 mM (Figure 3). A linear correlation also exists between copper ions and the GFP signal (r-squared value of 0.96), making it possible to quantify copper ions in a similar fashion to silver ions.

**Figure 3:**
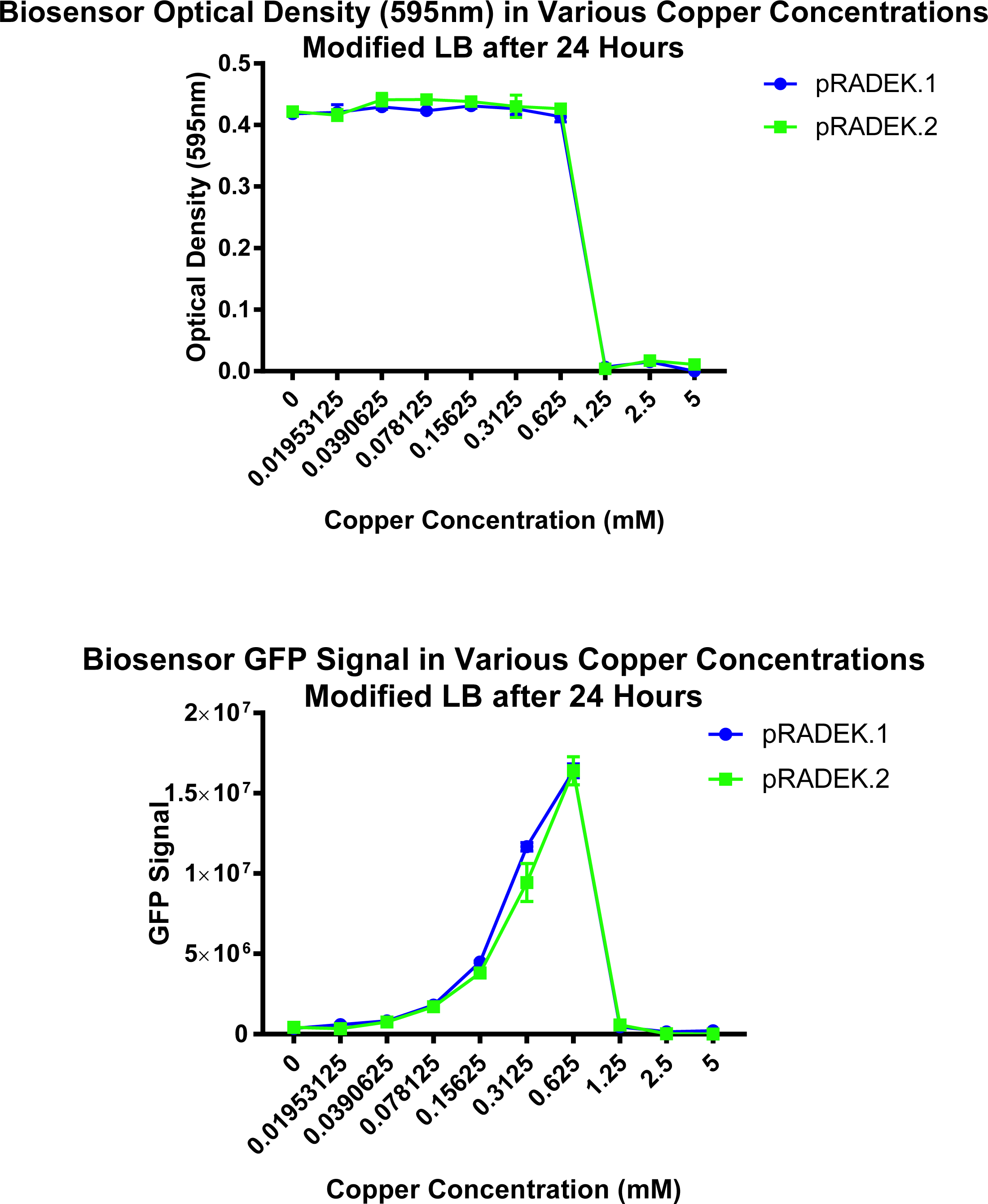
Line graph showing the optical density and GFP signal of biosensors RADEK.1 and RADEK.2 in modified LB at various silver concentrations.

### Discussion

Correlation between GFP signal and silver ion concentration makes it possible to quantify silver ions within a sample. The biosensor’s ability to correlate silver ions and GFP within a select range suggests a possible method for approximating the amount of silver ions within a sample. It was shown that the composition of the sample in which the biosensor is cultivated can affect the detection range and standard curve due to halides which affect the availability of silver ions. If a silver ion standard curve is made with each sample that needs quantification, this can yield a better approximation. In addition, since the correlations between silver and GFP are linear, it is possible to give a relative proportion of silver ion concentration between two samples. Although further testing is warranted to investigate the biosensor’s ability to detect different types of silver nanoparticles at various concentrations, results with research grade nanoparticles show that it is responsive at higher nanoparticle concentrations. Since nanoparticles release silver ions at a controlled rate over a period of time, it would make sense that a higher concentration is needed to obtain a signal from the biosensor. Theoretical estimations for the rate of silver ion release from nanoparticles are fairly complex and depend on various environmental factors and chemical properties of the nanoparticle. For example, the shape, size, and oxidative environment of the nanoparticle influences silver ion release rates. Stability and ion release rates of silver nanoparticles within different environments are important factors when determining potential applications. Since this biosensor is able to detect available silver ions released by the nanoparticle, determination of ion release rates and stability of silver nanoparticles may be another potential application for the biosensor. This is currently an ongoing area of work.

The biosensor was responsive to copper sulfide and exhibited an upward correlation trend with GFP and copper concentration similar to silver. This suggests that silver regulated GFP production via the *silRS* two-component system can also act as a mechanism for detecting copper(II) ions assuming the presence of a copper resistance gene. More interestingly, though, this result provides novel insight into the close homology of the copper and silver resistance mechanisms. Bioinformatic analysis of the two mechanisms encoded by *sil* and *cus* have confirmed many genetic similarities between various proteins. Rensing *et al.* found the silver ions activate the promoter regions found in the *cop* copper resistance mechanism (20). The findings from this study indicate the reverse; copper(II) ions can activate the *sil* promoter regions.

For long term environmental monitoring applications, plasmid-based engineering methods are likely not suitable due to factors such as copy number and plasmid loss. Further development can include integration of the *sil* operon fusions into the chromosome of *E. coli J53*. By doing this, the use of antibiotics for plasmid maintenance would not be required.

Since the biosensors were responsive to both copper sulfate and silver nitrate, methods for making the biosensor specific to one metal, or to differentiate between the two metals through their signal is warranted. One possible way would be to remove the copper resistance genes from pMG101 to determine whether the ability to detect the metal is affected, perhaps limiting detection to silver ions.

## Materials and Methods

### Molecular Cloning

Standard cloning protocols were used (21). These methods include calcium chloride transformations, restriction endonuclease digests and ligations, gel electrophoresis, PCR and gel extractions.

### Microbial Culturing and Competent Cells

Bacterial strains were either cultured in LB broth or on LB plates with a selection antibiotic as appropriate (Ampicillin 100 ug/ml or Kanamycin 25 ug/ml). Competent cells were made of both *E. coli* J53(pMG101), and *E. coli* DH5α using the method described by Sambrook (21). These cells were stored at -80°C until use.

### Plasmid and Genomic DNA Extractions and DNA Quantification

Plasmid DNA was extracted using anion exchange columns after an alkaline lysis. Genomic DNA was extracted with standard techniques: SDS lysis, chloroform extraction, and ethanol precipitation. Final DNA products were resuspended in a 50 ul solution, made of 0.2 M Tris at pH 8.0. Plasmid DNA was quantified using Nanodrop 2000 (22).

### PCR Amplification of Silver Regulated Promoters on pMG101

PCR primers were designed to amplify 244 and 245 bp of DNA before the transcription start point of both the *silE* and *silABC* respectively, according to pMG101 sequence at Genbank accession number PRJNA20285 (2). In addition, these primers contained EcoR1 and XbaI overhang regions. Although the promoter may have been less than 245 bp, the exact location of DNA binding protein sites have not been characterized yet; 245 bp was safely large enough when comparing with defined promoters on other similarly regulated heavy metal resistance mechanisms like the cus operon. Refer to Table 1 for primer sequences. PCR amplification was done using the KOD amplification system, and a touchdown cycle program. Amplification was confirmed by running products on a 2% agarose gel.

**Table 1:**
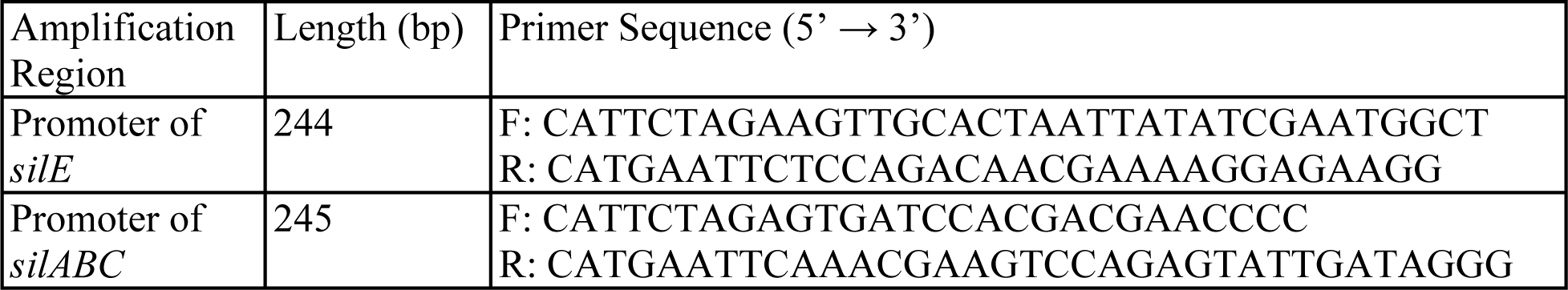
Primers for the amplification of the *sil* operon promoters

### Engineering Vector Construct

pFPV25.1, containing a fluorescent mutGFP gene (23), was purchased from Addgene (catalog number 20668). The plasmid contained an ampicillin resistance marker which pMG101 happens to also convey resistance to. Thus, the fluorescent GFP gene from pFPV25.1 was cleaved using restriction enzymes (EcoR1 and HindIII) and ligated with EcoR1 HindIII cleaved pK19mobsacB (24). Ligation products were recovered by transformation of *E. coli* DH5α (25), followed by selection on ampicillin and kanamycin. The insertion of the GFP gene into the plasmid pK19mobsacB was confirmed through restriction enzyme digestion. The newly constructed plasmid was designated pK19mobsacGFP.

Next, the promoter controlling the production of GFP in pK19mobsacGFP was replaced with the silver regulated promoter found in pMG101. The current promoter region from pK19mobsacGFP was taken out and replaced with the PCR amplified promoter region (from *silE* being 244 bp in length, and from *silABC* 245bp in length) using restriction enzymes EcoRI and XbaI. The resulting two plasmids pRADEK.1 (containing *silE* promoter) and pRADEK.2 (containing *silABC* promoter) were obtained by transformation of *E. coli* DH5*α*. Successful promoter fusions were confirmed with digestion using the same restriction enzymes. pRADEK.1 and pRADEK.2, then transformed into *E. coli* J53(pMG101) and selected using kanamycin and ampicillin. The resulting biosensor strains were subjected to further experimentation.

### Biosensor Sensitivity Assay to Silver Ions and Copper Ions

The ability of the constructed biosensor strains to detect silver nitrate and copper sulfide was tested in a 96-well microtiter plate by using the 2-fold microdilution method with slight modifications. The biosensor was pre-cultivated in LB with 100 ug/ml Amp and 25 ug/ml Kan overnight at 37°C (200 rpm), and then diluted to obtain an optical density reading of 0.2 at 600 nm. Ionic silver, and copper concentrations were tested at 10 mM, followed by 2-fold concentration dilutions until 0.02 mM. Control wells contained the bacterium in LB alone, LB and silver nitrate concentrations alone, and LB alone. After 24 h of incubation at 37°C shaking at 200 rpm, the GFP light signal and optical density (595 nm) were measured. A Molecular Devices (Sunnyvale, California) FilterMax F5 was used to read standard clear well microplates. Before measurements were taken, the plate was shaken for 5 seconds. GFP fluorescence was measured using bandpass filters centered on 485 nm for excitation, and 535 nm for emission. This test was repeated for two types of LB media, one being the standard recipe, and second one being modified to contain no sodium chloride (referred to as modified LB). Sodium chloride can affect the availability of silver ions through the formation of a silver chloride precipitate. By removing sodium chloride from the LB media, a better measurement of the biosensors ability to detect free silver ions can be achieved. This test was repeated at various start concentrations to obtain data points in ranges that the biosensor was able to detect. In addition, the GFP signal for each concentration was subtracted from LB and silver to remove background noise. Data was examined for a positive correlation (linear correlation coefficient) between silver, and copper concentration and GFP signal.

*E. coli* J53(pMG101), *E. coli* J53 (pK19mobsacGFP), and silver sensitive *E. coli DH5α* were included as controls.

### Biosensor Sensitivity Assay to Silver Nanoparticles

Research grade silver nanoparticles were graciously made and donated by Dr. Kitaev’s Lab at Wilfrid Laurier University. Two types of nanoparticles, provided at a stock concentration of 0.13 mM, each suspected to have different silver ion release rates, were tested in this experiment. One silver nanoparticle type was between 10-15 nm and contained a mix of particle shapes, with the other being around 65 nm and decahedra (pentagonal prism) in shape (26, 27). Nanoparticles were concentrated using vacuum-centrifugation until 0.26 mM. A similar assay to silver and copper was used expect with the modification that silver concentrations only started at 0.1 mM. This was due to the limited amount of available silver nanoparticles.

## Acknowledgments

This work was supported by an NSERC Discovery Grant. J.R.H was a Mitacs Elevate Fellowship recipient.

We are grateful to Dr. Alex O’Neill of University of Leeds for the generous gift of strain J53(pMG101). We would like to thank Dr. Vladimir Kitaev at Wilfrid Laurier University for graciously donating silver nanoparticles for this study. In addition, we are grateful to Dr. Marc Habash at the University of Guelph for discussions and suggestions pertaining to the development of the biosensor.

